# Graph analytics for phenome-genome associations inference

**DOI:** 10.1101/682229

**Authors:** Davide Cirillo, Dario Garcia-Gasulla, Ulises Cortés, Alfonso Valencia

**Affiliations:** Barcelona Supercomputing Center (BSC), C/ Jordi Girona 29, 08034, Barcelona, Spain; ICREA, Pg. Lluís Companys 23, 08010, Barcelona, Spain; Universitat Politècnica de Catalunya (UPC), C/ Jordi Girona 31, 08034, Barcelona, Spain

## Abstract

**Motivation:** Biological ontologies, such as the Human Phenotype Ontology (HPO) and the Gene Ontology (GO), are extensively used in biomedical research to find enrichment in the annotations of specific gene sets. However, the interpretation of the encoded information would greatly benefit from methods that effectively interoperate between multiple ontologies providing molecular details of disease-related features.

**Results:** In this work, we present a statistical framework based on graph theory to infer direct associations between HPO and GO terms that do not share co-annotated genes. The method enables to map genotypic features to phenotypic features thus providing a valid tool for bridging functional and pathological annotations. We validated the results by (a) supporting evidence of known drug-target associations (PanDrugs), protein-protein physical and functional interactions (BioGRID and STRING), and common pathways (Reactome); (b) comparing relationships inferred from early ontology releases with knowledge contained in the latest versions.

**Applications:** We applied our method to improve the interpretation of molecular processes involved in pathological conditions, illustrating the applicability of our predictions with a number of biological examples. In particular, we applied our method to expand the list of relevant genes from standard functional enrichment analysis of high-throughput experimental results in the context of comorbidities between Alzheimer’s disease, Lung Cancer and Glioblastoma. Moreover, we analyzed pathways linked to predicted phenotype-genotype associations getting insights into the molecular actors of cellular senescence in Proteus syndrome.

**Availability:** https://github.com/dariogarcia/phenotype-genotype_graph_characterization

## Introduction

The phenome can be defined as the totality of all distinct variants of phenotypic characteristics (traits) expressed by a cell, a tissue, an organ, an organism, or a species, under the influence of both genetic variation and environmental factors (Mahner and Kary. J Theor Biol. 1997). Finding associations between phenome, genome and environment is of utmost priority in biomedicine as it could lead to the identification of the molecular drivers underlying human traits and diseases. Genome-wide association studies (GWAS) have been extensively carried out to dissect complex traits (Beck et al. E J Hum Genet. 2014). Moreover, reverse genetic approaches such as phenome-wide association studies (PheWAS) (Bush et al. Nat Rev Genet. 2016), as well as large systems genetics infrastructures (Li et al. Cell Syst. 2018), have been recently developed. Nonetheless, the phenome-genome association is hampered by biological complexity (Hall et al. Trends Genet. 2016) and lack of consensus on the evidence level required to establish pathogenicity and disease susceptibility (Strande et al. Am J Hum Genet. 2017). Yielding a definitive molecular diagnosis from the genotype or a confident clinical diagnosis from the phenotype is arduous, especially in absence of richly genotyped or phenotyped sets like the case of rare diseases (Wright et al. Nat Rev Genet. 2018). Categorical approaches use biomedical ontologies to systematically capture genotypic and phenotypic attributes to facilitate the identification of discriminative patterns of molecular and clinical features. An ontology is a domain-specific knowledge formalization based on sets of entities with relations operating among them (Schulze-Kremer Pac Symp Biocomput. 1998). Biological ontologies are recognized as essential in the grand challenges of biomedical research (Hoehndorf et al. Brief Bioinform. 2015), and community efforts like the Critical Assessment of Functional Annotation (CAFA; https://biofunctionprediction.org/cafa/) have been created.

In this work, we developed and evaluated a method based on graph theory to infer phenome-genome associations by using the Human Phenotype Ontology (HPO) and the Gene Ontology (GO). **To our knowledge, this is the first method developed specifically to infer associations between HPO and GO terms that do not exhibit any already existing co-annotation.** Indeed, tools designed to address ontology interoperability generally rely on the availability of co-annotations between specific terms, such as HPO2GO (Doğan PeerJ 2018), which annotates a gene to a certain HPO term based on supplemental co-annotated GO terms, and Phevor (Singleton et al. Am J Hum Genet. 2014), which identifies distinct annotations relevant for a HPO-GO co-annotation of interest. The applications of our method show that our predictions improve the molecular interpretation of distinct pathological processes. In particular, we enhanced standard functional enrichment analysis of genes with expression patterns consistent with observed comorbidity between Alzheimer’s disease, Lung Cancer, and Glioblastoma. Moreover, we uncovered relevant molecular insights for the rare condition known as Proteus syndrome.

## Methods

### Ontologies and annotations

For our experiments, we used the Gene Ontology (GO) (Ashburner et al. Nat Genet. 2000) and the Human Phenotype Ontology (HPO) (Köhler et al. Nucleic Acids Res. 2017), available at OBO Foundry Permanent URLs (PURLs) http://purl.obolibrary.org/obo/. We used GO ontologies released on 29 June 2013 and 23 May 2017, and HPO ontologies released on 31 May 2013 and 13 April 2017 (Suppl. Table 1). We only considered human GO annotations associated with experimental evidence codes. As for the gene annotations, we used the human GO Annotation (GOA) for UniProt version 117 (released on 05 March 2013) and version 168 (released on 09 May 2017), available at https://www.ebi.ac.uk/GOA, and the gene-to-phenotype associations (“ALL_SOURCES_ALL_FREQUENCIES”) released on 01 July 2013 and 29 June 2017, available at https://github.com/Phenomics/HPO-archive.

### Model implementation

Our model assumes the existence of two ontologies, P and G, a set of genes H, and two sets of gene annotations <p,h> and <g,h> where p∈P, g∈G and h∈H. The objective of the model is to, given a pair of terms <p,g> which do not have shared annotated genes, estimate the likelihood that such an annotation should exist. For this purpose, we separated all possible pairs <p,g> in two sets: those which have a shared annotated gene h (henceforth called connected pairs), and those which do not (henceforth called disconnected pairs). We then built a representation of each of those two sets, such that when a given pair <p’,g’> is provided we can estimate the likelihood of the pair to belong to the set of connected pairs and to the set of disconnected pairs. To build these representations we created a composed graph, which includes all terms from both ontologies, all “is-a” relations between terms of the same ontology, and all genes annotated to the ontologies (Figure 1).

**Figure 1.**
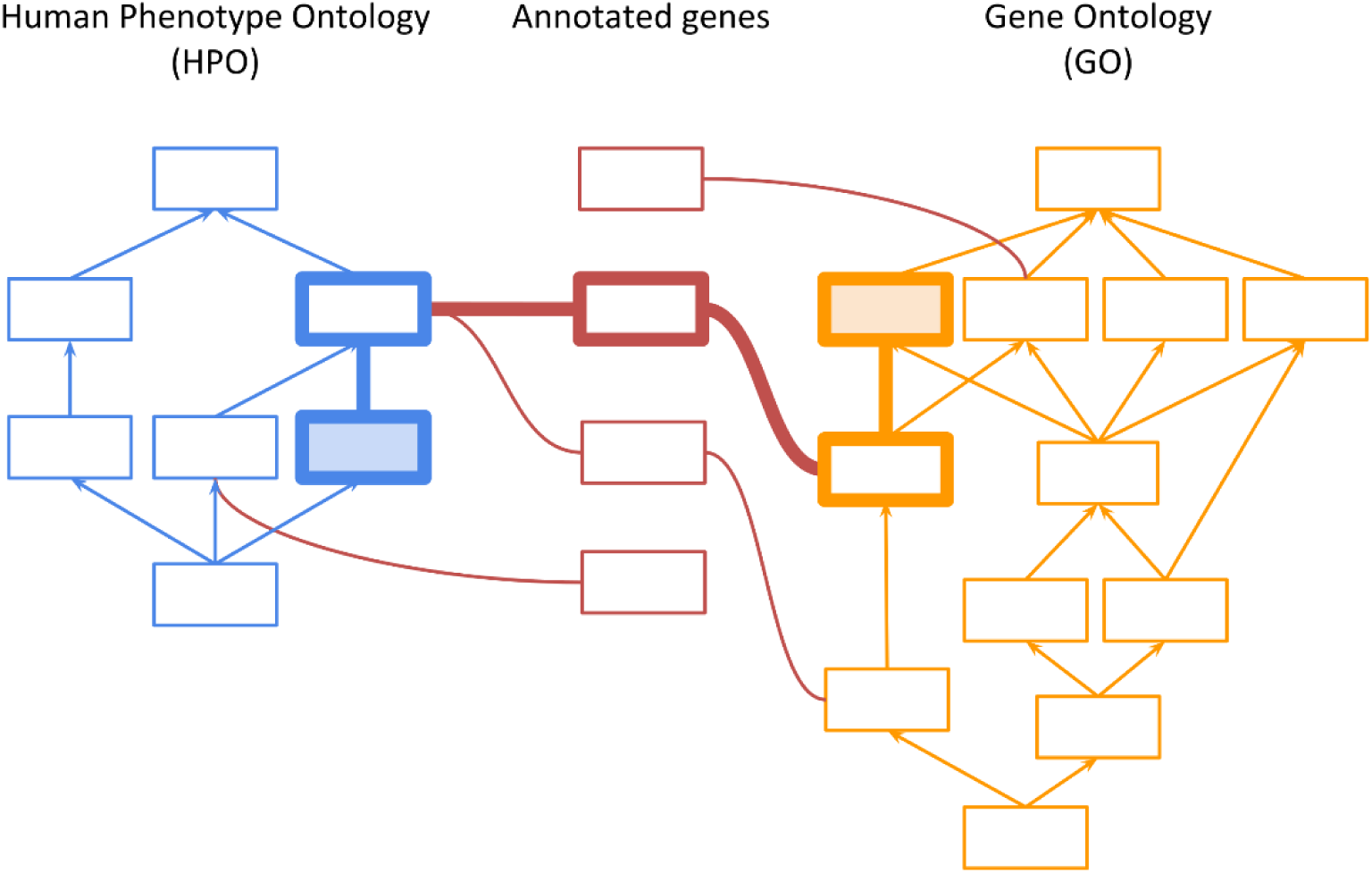
Illustration of the composed graph used for inference. Human Phenotype Ontology (HPO) (in blue) and Gene Ontology (GO) (in orange) consist of vertices (terms) and edges (“is_a” relationships). Genes with at least one annotation to an ontology term (in red) are added to the graph, representing a connection layer between the two ontologies. Thick lines highlight an example path connecting an HPO term (blue filled box) and a GO term (orange filled box) that do not directly share a co-annotated gene.

### Representations of connected and disconnected pairs

The representations of the two sets of connected and disconnected pairs are generated after random sampling. For each randomly sampled connected and disconnected pair <p,g>, we computed the total number of paths that lead from one term to the other, with a maximum distance of 4 edges, chosen to limit both the combinatorial explosion (O(*n*!) for a graph with *n* nodes) and paths’ sparsity (88 possible paths with a maximum of 4 steps for a given pair <p,g>). Paths are walked through the composed graph in a two-way unconstrained manner to allow for a deeper exploration of the graph structure which is crucial for inference (Mohamed et al. Lecture Notes in Computer Science. 2017).

Along with the total number of paths, we also computed the type of path, which is defined by the type of vertices being visited at each hop (p for vertices in P, h for vertices in H, and g for vertices in G). For example, a path may start in the source HPO term p^1^, which is connected with a gene h^1^, which is connected with another HPO term p^2^, which is connected with another gene h^2^, which is finally connected to the target GO term g^1^. Such a path would be of type “phphg”. The random sampling continued until stabilization of path statistics is attained (Suppl. Table 2). In particular, using 2013 ontology releases, we randomly sampled 123,834 disconnected pairs (0.3% of the total disconnected pairs) and 34,499 connected pairs (10% of the total connected pairs), and computed the average number of paths per path type, and its standard deviation. Same statistics have been computed for 2017 ontology releases, from which we randomly sampled 111,952 disconnected pairs (0.2% of the total disconnected pairs) and 38,703 connected pairs (5% of the total connected pairs).

### Likelihood estimation

Once we generated the representations of the randomly sampled connected and disconnected pairs, we assessed the likelihood of a given pair <p’,g’> to belong to each of those sets. To do so, we measured how well the number and type of paths between <p’,g’> fit the distributions of paths within the sets of connected and disconnected pairs. In particular, we estimated the probability density function (PDF) per path type, assuming a normal distribution. The likelihood of the given pair to belong to a set is then computed as the product of all its PDFs:

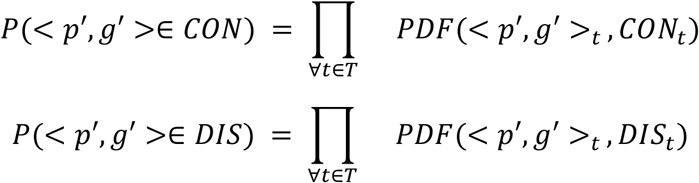

where CON is the representation of connected pairs, DIS is the representation of disconnected pairs, <p’,g’> is any pair of terms not used to build CON or DIS, T is the set of all path types, <p’,g’>t is the number of paths between p and g of type t, and CONt is the mean and standard deviation of paths of type t within CON (analogous for DIS).

For any given pair <p’,g’>, this process generates a likelihood of the pair being connected and a likelihood of the pair being disconnected. We compute those likelihoods for a large number of random disconnected pairs, ensuring that the ones used to generate the representations are not considered. We generated 1,000,682 pairs (of a total of 45 million) by using 2013 ontology releases, and 244,315 pairs (of a total of 82 million) by using 2017 ontology releases. In order to obtain the most reliable candidate pairs to be connected or disconnected, likelihoods have been sorted as follows:

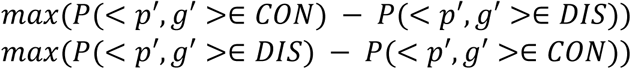

We then kept the top 2,000 pairs resulting from each sorting, and used these as our ranked predictions (henceforth called candidate pairs and non-candidate pairs, respectively).

### Evaluation metrics

Method performance has been evaluated by computing the Area under Receiving Operator Characteristic Curve (AUROC), which measures the probability that positive instances are ranked higher than negative ones. An HPO-GO association predicted using the 2013 ontology releases is a true positive (TP) if that association occurs in the 2017 ontology releases (i.e. at least one gene is co-annotated). If no association occurs, a positive prediction is a false positive (FP). If the algorithm fails to predict a true association, it is a false negative (FN). If no true association occurs, a negative prediction is a true negative (TN). We also computed established metrics for the goodness of fit of prediction (precision, accuracy, F1-score, Matthews Correlation Coefficient (MCC)) after identifying a decision threshold (optimal cut-off point) from the ROC curve by maximizing the Youden’s index, i.e. the difference between recall and False Positive Rate (FPR). All the analyses have been performed using R package pROC.

To compare 2013 and 2017 ontology releases, we split the gene annotations into three categories: annotations that are stable across the releases; annotations that are new in the more recent release; annotations that changed from one release to the other. Given the genes annotated to a predicted HPO-GO terms pair, we measured the intersection between stable annotations of one ontology and new annotations of the other (Suppl. Figure 3).

### External databases

Several databases covering different types of supporting evidence of our predictions were employed. PanDrugs (Perales-Patón et al. Public Health Genomics. 2017) (build of April 2017) is a computational method to prioritize therapies by considering the biological relevance of altered genes in cancer, their therapeutic vulnerability and drugs clinical application. In our analysis, we queried PanDrugs for genes, FDA approved drugs, pathways the gene annotates to, and pathologies the drug is prescribed for. STRING (Szklarczyk et al. Nucleic Acids Res. 2017) is a database of physical and functional interactions. We utilized STRING v10.5 human protein network data. The Biological General Repository for Interaction Datasets (BioGRID) (Chatr-Aryamontri et al. Nucleic Acids Res. 2017) (built 3.4.152, September 2017) is a database of protein, genetic and chemical interactions. We utilized human physical protein-protein interactions experimentally verified by co-localization, co-purification, FRET, and two-hybrid assays (Cirillo et al. Nucleic Acids Res. 2015). Network analysis was performed using R package igraph. Reactome (Fabregat et al. Nucleic Acids Res. 2018) is a curated database of human-specific pathways. We performed a Reactome pathway enrichment analysis (Benjamini-Hochberg adjusted p-value cutoff 0.01) using the R package ReactomePA.

### Additional analyses

Information content (IC), or Shannon information has been computed as in Alterovitz et al. Nucleic Acids Res. 2007. Go term enrichment analysis has been performed using R package GOstats with Dunn-Šidák multiple conditional test correction at 5%. Disease matching has been performed using PhenoGrid (Smedley et al. Database 2013), a tool for visualizing semantic similarity between phenotypes and a target group. We used “Homo sapiens (diseases)” as target group, and reported the disease with the highest score and the highest number of matches.

## Results

### Performance evaluation

We generated predictions using the 2013 releases of HPO and GO ontologies and validate them using the 2017 releases (Methods). In particular, our predictions consist of a set of 2000 candidate pairs (i.e. HPO-GO term pairs predicted to be co-annotated) and 2000 non-candidate pairs (i.e. HPO-GO term pairs predicted not to be co-annotated) (Suppl. Table 3). As 65 candidate pairs and 111 non-candidate pairs do not have experimentally validated gene annotations in the 2017 ontology releases, we excluded them from the performance analysis, and balanced the two classes based on the least populated one, resulting in 1889 candidate pairs and 1889 non-candidate pairs.

327 candidate pairs (17%) and 1882 non-candidate pairs (99.6%) are correctly found to be co-annotated and not co-annotated, respectively, in 2017 ontology releases. Only 7 non-candidate pairs (0.4%) are co-annotated in 2017 ontology releases, while the remaining 1562 candidate pairs (83%) are not co-annotated, as expected due to the considerable fraction of genes still missing a functional characterization (Cozzetto and Jones, Methods Mol Biol. 2017). Morover, the average number of annotated genes is higher in candidate pairs compared to pairs predicted not to be co-annotated (Suppl. Figure 1).

For performance evaluation, we measured the Area Under the Receiver-Operating Characteristic (AUROC) curve (0.77; p-value < 1e-03 after label shuffling; Figure 2). By bootstrapping the AUROC score, we computed that the performance has a 95% confidence interval between 0.75 and 0.79 (Suppl. Figure 2). Other performance assessment metrics for binary classification have also been evaluated (precision 0.92, recall 0.59, F1-score 0.67, MCC 0.30) (Methods).

**Figure 2.**
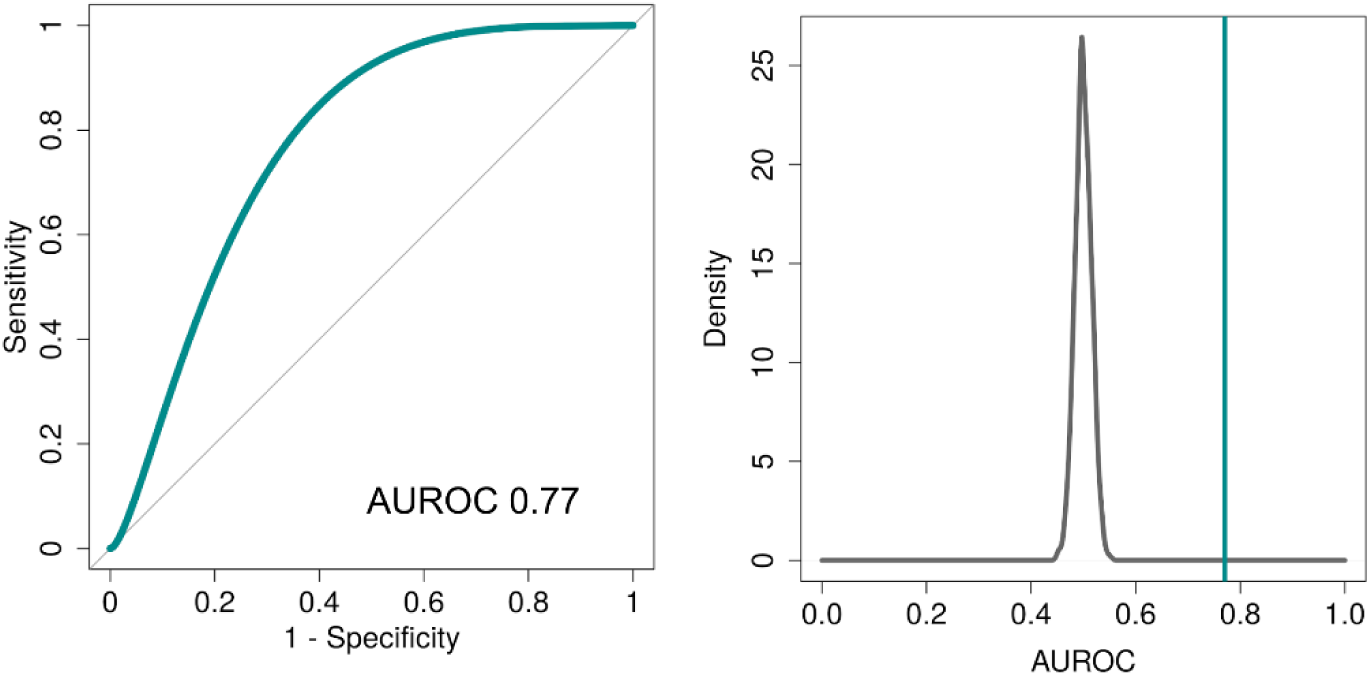
Performance evaluation of predictions using 2013 ontology releases (Methods). The corresponding Area Under the Receiver-Operating Characteristic (AUROC) is 0.77 (p-value 1e-03 after label randomization, in grey, right panel; binomial smoothing).

### Accumulation of gene annotations over time

The rate of accumulation of ontology annotations across time is a steady process leading to a continuous reshaping and updating in the dedicated repositories (Tomczak et al. Sci Rep. 2018). For instance, by comparing 2013 and 2017 ontology releases we found that 48% of experimentally validated GO annotations were maintained while 45% were new and 7% were dropped. In the case of HPO annotations, 48% of those were the same in the two releases, 41% were new, and 11% were dropped (Figure 3).

**Figure 3.**
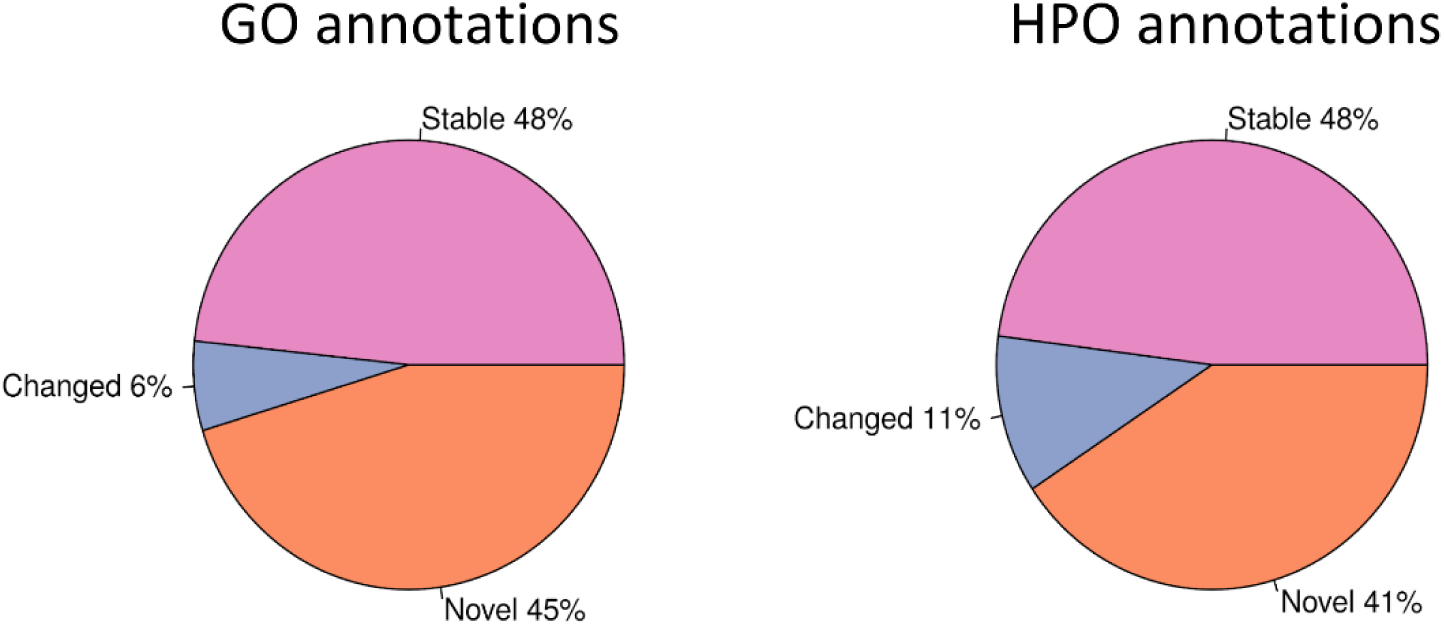
Proportion of GO and HPO annotations that have been maintained (stable; light purple), dropped (changed; dark purple), and added (novel; orange) from 2013 to 2017 ontology releases.

We sought to evaluate our HPO-GO predicted associations in terms of number of annotations accumulated over time between the two ontologies (Methods; Suppl. Figure 3). In particular, given a predicted HPO-GO terms pair, we counted how many genes annotating GO between 2013 and 2017 are the same genes of maintained or acquired annotations of HPO, and vice-versa. **We found that 81 out of the 327 correctly predicted candidate pairs (24.8%) have gene annotations (n = 102) that are new in both HPO and GO in the 2017 releases. 110 out of the 327 pairs (33.6%) have gene annotations (n = 146) that are new in GO and maintained in HPO since 2013. Finally, 172 out the 327 pairs (52.6%) have gene annotations (n = 200) that are new in HPO and maintained in GO**. This analysis shows that our predictive model is able to overcome the differences in the process of annotation accumulation in the two ontologies (Tomczak et al. Sci Rep. 2018).

### Ontology levels and information content

We compared graph levels and Information content (IC) (Methods) of the ontologies and our predictions. No relevant differences in the distribution of graph levels between the ontologies and our predictions can be observed (Suppl. Figure 4A) meaning that our predictions are not biased towards specific areas of the ontologies from which are drawn. Conversely, the distribution of IC of the candidate pairs differs drastically from that of the ontologies (Suppl. Figure 4B) showing that the specificity of our predictions is not prejudiced by the difference in abundance between highly specific and more general terms in the ontology trees. In summary, our predictions span all ontology levels and are enriched in terms with intermediate annotation specificity making them suitable for a broad range of applications.

### Supporting evidence in external databases

In order to assess the quality of the predictions, we employed several databases covering different types of supporting evidence for our candidate pairs. PanDrugs (Methods) is a platform devoted to prioritize anticancer drug treatments by integrating data on drug-target associations. We found that 2728 genes annotated to predicted HPO-GO pairs are target of 3503 approved drugs in 98 pathologies affecting 335 pathways. We observed that the co-occurrence of genes annotated to candidate pairs is higher than those predicted not to be co-annotated (p-value < 2.2e-16, Kolmogorov-Smirnov test; Figure 4).

**Figure 4.**
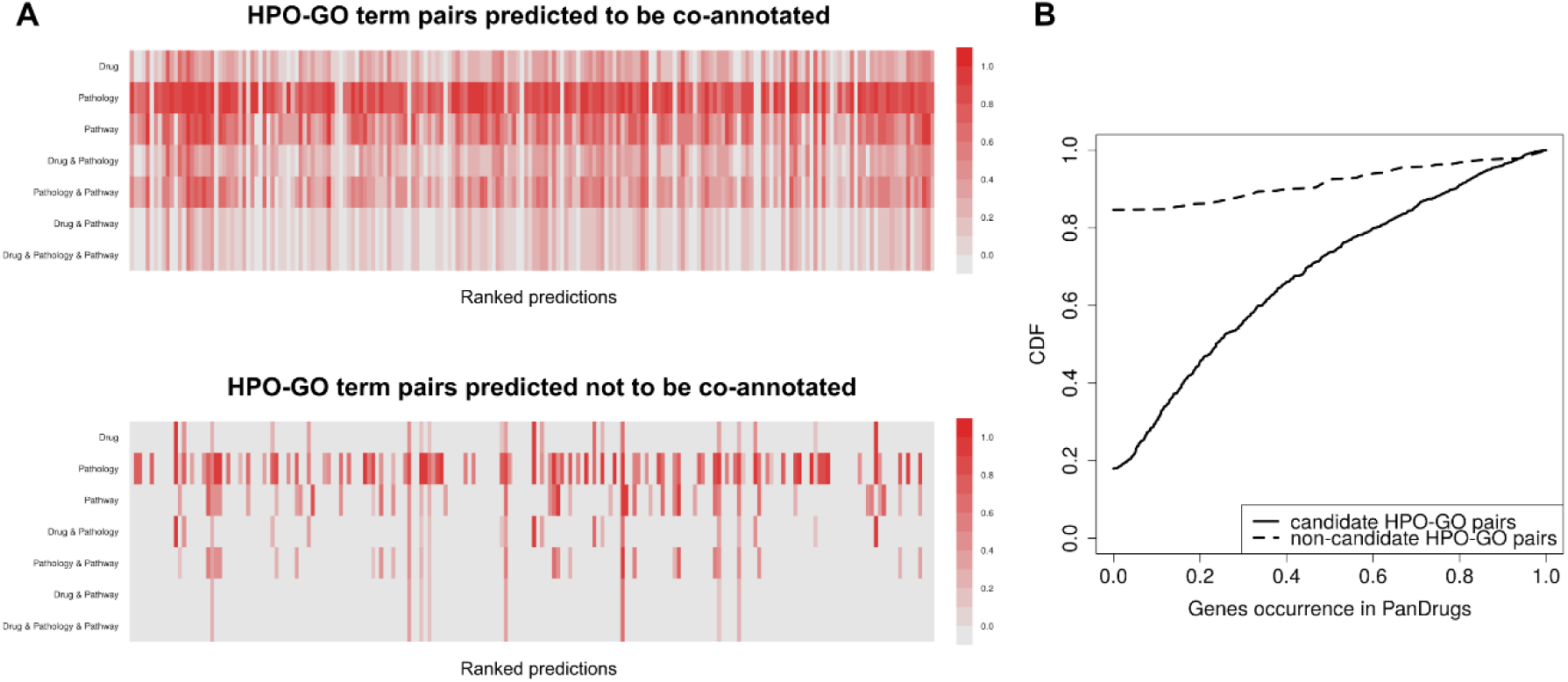
[A] Co-occurrence of genes annotated to predicted HPO-GO terms pairs in PanDrugs entries (Drugs, Pathology, Pathway, and combinations). Top and bottom 10% (200 predicted pairs) are shown in the two panels, respectively, with increasing prediction ranking from left to right. Colors (gray to red) indicate the number of co-occurrencies normalized by the total number of co-occurrencies for a given PanDrug entry. [B] Cumulative Distribution Function (CDF) of normalized co-occurrences. A significant difference between the two types of predicted HP-GO term pairs is observed (p-value < 2.2e-16, Kolmogorov-Smirnov test; >80% of non-candidate pairs are not found in PanDrugs).

We found evidence for products of genes annotated to predicted HPO-GO pairs to be physical interactors reported in BioGRID (Methods). BioGRID is a curated database of interaction data. Interactors annotated to candidate pairs create a network of 1693 nodes, 2363 edges, and 93 connected components (one of 1480 nodes, the others of 2.3 nodes on average). Interactors annotated to non-candidate pairs, instead, create a considerably smaller network of 134 nodes, 86 edges, and 48 connected components (one of 12 nodes, the others of 2.6 nodes on average). Moreover, closeness centrality of the largest components of candidate pairs is significantly higher than the one of non-candidate pairs (p-value 1.161e-09, one-sided Wilcoxon signed-rank test).

Products of genes annotated to predicted HPO-GO pairs are also found in STRING (Methods), an integrative database of functional interactions. In particular, we found 7958 genes annotated to candidate pairs involved in 212936 unique pairwise interactions, and 3301 genes annotated to non-candidate pairs involved in 9700 unique pairwise interactions. A significant difference between the fraction of co-occurrence of genes annotated to candidate pairs and that of genes annotated to non-candidate pairs is observed at different confidence score cutoffs (p-value 3.289e-67 at low confidence; p-value 1.251e-290, at high confidence; two-sided Wilcoxon signed-rank test; Suppl. Figure 5).

### Improving the interpretation of standard GO enrichment analysis

We used our predictions to improve the interpretability of standard enrichment analysis on gene sets, such as the widely used GO enrichment analysis. GO enrichment analysis is a common approach for the interpretation of gene expression data as it allows the identification of over-represented functional categories in arbitrary sets of genes. In this application, we analyzed six sets of genes that are up- (Up) or down-regulated (Down) in three pathologies, Alzheimer’s disease (AD), Lung cancer (LC) and Glioblastoma (GBM), showing distinct expression patterns consistent with direct and inverse comorbidities observed at epidemiological level (Sánchez-Valle et al. Sci Rep. 2017). For each of the six gene sets, we identified over-represented GO terms (Methods), obtaining a total of 244 functional categories overlapping with our candidate pairs. We found that the genes annotated to the HPO terms that are predicted to associate with the enriched GO terms can be used to expand the list of relevant genes from the high-throughput experimental results (Suppl. Figure 6A), providing additional information to the molecular characteristics of gene sets. The average number of additional genes recovered through our approach is 18.19 (Suppl. Figure 6B), being genes of LC sets associated with more abundantly annotated HPO terms. Remarkably, we found that the overlap of the recovered genes among the three pathologies recapitulates the results from the original study (Suppl. Figure 6C) suggesting that deregulated genes determining specific phenotypes are shared among diseases with comorbidity relations. For instance, 22 genes are annotated to HPO terms predicted to be associated to GO terms that are enriched in AD.Up and LC.Down. The gene ANG is annotated, among other phenotypes, to emotional lability (HP:0000712) and agitation (HP:0000713) (Moretti et al. Expert Rev Neurother. 2006), predicted to be associated with immune response (GO:0006955) and lipid binding (GO:0008289), functions enriched in both AD.Up and LC.Down. The gene SLC7A7, instead, is annotated to leukopenia (HP:0001882), predicted to associate with response to growth factor (GO:0070848), enriched in AD.Up, and, among others, splenomegaly (HP:0001744), associated to transmembrane receptor protein tyrosine kinase signaling pathway (GO:0007169), enriched in LC.Down. Spleen metastasis are indeed rare in LC (Iguchi et al. Exp Ther Med. 2015), while growth factors-based therapies are at the forefront of AD treatment (Duncan and Valenzuela Stem Cell Res Ther. 2017). Those results show the potential of mapping relevant phenotypic features to enriched molecular functions in order to facilitate gene candidate prioritization and treatment design.

### Pathway analysis of candidate pairs

Candidate pairs entail 227 distinct clusters: 226 ranging from 2 to 38 terms in size and one of 1269 connected terms (Suppl. Figure 7). For each cluster, we performed a pathway enrichment analysis after merging the genes annotated to HPO and GO terms and excluding pathways that are over-represented in HPO- and GO-annotated genes separately. We found that 100 clusters out of 227 show enrichment in processes linking specific phenotypes to gene functional categories. For instance, the pool of genes PIK3R1, WAS, HLA-DPB1, annotated to the respiratory inflammation sinusitis (HP:0000246), and BCL10, RIPK2, UBB, annotated to protein ubiquitination (GO:0031398), are enriched in T cell receptor (TCR) signaling (Reactome ID 202403) (p-value 0.00025), confirming the role of this protein modification mechanism in inflammatory diseases and recurrent infections (Hu and Sun Cell Res. 2016). More complex associations emerge from the analysis of bigger clusters of candidate pairs. For instance, oxidative stress induced senescence (Reactome ID 2559580) (p-value 0.0027) is enriched in genes CDKN2C, CDKN2B, annotated to hypocholesterolemia (HP:0003146), hand oligodactyly (HP:0001180), posterior subcapsular cataract (HP:0007787), ileus (HP:0002595), retinal nonattachment (HP:0007899), and increased urinary cortisol level (HP:0012030), together with 19 genes annotated to RNA binding (GO:0003723), cadherin binding (GO:0045296), and nuclear body (GO:0016604) (Suppl. Figure 7). According to PhenoGrid (Smedley et al. Database 2013), those symptoms are consistent with Proteus syndrome (Monarch ID MONDO:0008318), a rare disease characterized by progressive overgrowth of the skeleton, skin, and adipose tissues. Importantly, this syndrome has been associated with a variant in the gene AKT1 whose deregulation promotes senescence events (Skeen et al. Cancer Cell 2006).

## Discussion

The use of biological ontologies has grown drastically in the last decades. Nowadays, they are recognized as crucial resources for the most important challenges in genomics research, especially for rare diseases. In this work, we developed a statistical framework based on graph theory to infer direct associations between HPO and GO terms. The method enables a direct mapping between phenotypic features (HPO terms) and biological processes, molecular functions, and cellular components (GO terms). The statistical approach is based on topological features of the ontology trees that are used to infer the likelihood of association between terms devoid of a direct cross-reference. **The method achieved high performances (AUROC 0.75-0.79 with 95% confidence interval) in a retrospective benchmark in which predictions of 2013 ontology releases are evaluated using ontologies from 2017.** By comparing stable and new annotations of those releases, we showed that our predictions recapitulate a large fraction of annotations accumulated over time between 2013 and 2017. Furthermore, we generated predictions of 2017 ontology releases and found that our findings are supported by evidence deposited in several databases (PanDrugs, BioGRID, STRING, Reactome), namely drug-target associations, protein-protein physical and functional interactions, and common pathways. **We predicted phenome-genome associations for specific biological cases showing that our predictions improve the molecular interpretation of distinct pathological processes. In particular, we applied our method to the analysis of large-scale comorbidity studies expanding the results of standard GO term enrichment analysis to gain deeper insights into the pathological processes.** We found that deregulated genes in AD, LC and GBM are associated to specific phenotypic annotations showing expression patterns that are consistent with observed direct and inverse comorbidity relations among those conditions.

We showed that groups of predicted HPO-GO associations encapsulate relevant information about genes involved in specific pathologies, namely inflammatory diseases and rare diseases. We identified molecular players putatively acting in Proteus syndrome. Interestingly, RNA-binding emerged as a relevant functional category, corroborating the importance of this molecular activity in cell-growth and proliferation. In conclusion, we introduced a method to infer pairwise associations between HPO and GO terms that can be used to identify symptoms-gene function associations, thus enabling ways to refine results from standard functional analysis of high-throughput experimental results.

Finally, it is important to highlight that the algorithm can be adapted to any arbitrary pair of ontologies, making it a powerful statistical framework to integrate heterogeneous source of biological knowledge.

## Acknowledgments

The authors thank Javier Béjar and Antonio García Díaz for stimulating discussions.

