## Supplementary Material for "Graph analytics for phenome-genome associations inference"

### Supplementary Figures

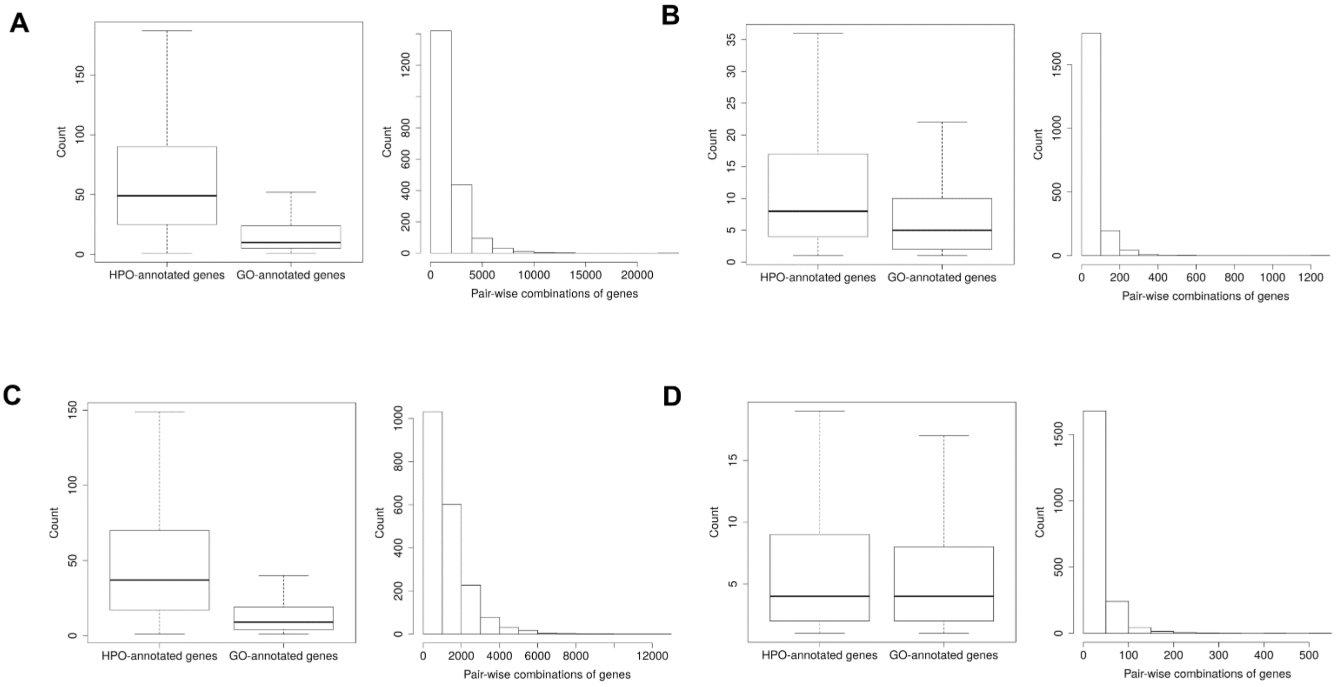


**Supplementary Figure 1.** Genes annotated to predicted HPO-GO pairs (see Suppl. Table 1) and distribution of pair-wise gene combinations. [A] (left panel) Count of genes annotated to HPO-GO terms pairs predicted to be co-annotated (2017 ontology releases). HPO-annotated genes: tot 3383, min 1, max 1945, median 49, average 80.40, SD 123.51. GO-annotated genes: tot 10527, min 1, max 2738, median 10, average 29.55, SD 110.46. (right panel) The total number of gene combinations is 3454753; the average number of gene combinations is 1727.37. [B] (left panel) Count of genes annotated to HPO-GO terms pairs predicted not to be co-annotated (2017 ontology releases). HPO-annotated genes: tot 2884, min 1, max 231, median 8, average 12.93, SD 15.32. GO-annotated genes: tot 5677, min 1, max 217, median 5, average 8.45, SD 12.31. (right panel) The total number of gene combinations is 116165; the average number of gene combinations is 58.08. [C] (left panel) Count of genes annotated to HPO-GO terms pairs predicted to be co-annotated (2013 ontology releases). HPO-annotated genes: tot 2672, min 1, max 1524, median 37, average 60.58, SD 95.49. GO-annotated genes: tot 7385, min 1, max 2413, median 9, average 29.69, SD 128.76. (right panel) The total number of gene combinations is 2611270; the average number of gene combinations is 1306.28. [D] (left panel) Count of genes annotated to HPO-GO terms pairs predicted not to be co-annotated (2013 ontology releases). HPO-annotated genes: tot 2050, min 1, max 78, median 4, average 7.48, SD 9.42. GO-annotated genes: tot 4013, min 1, max 106, median 4, average 6.84, SD 9.26. (right panel) The total number of gene combinations is 60252; the average number of gene combinations is 30.27.

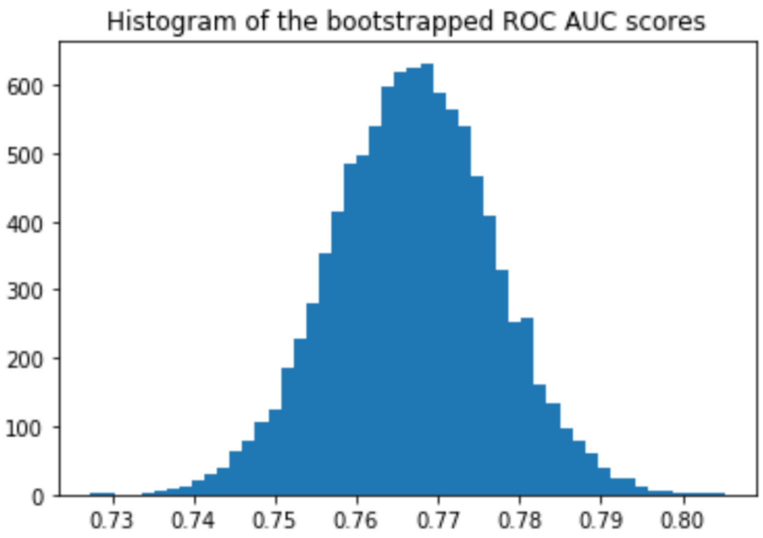


**Supplementary Figure 2.** Histogram of the bootstrapped ROC AUC scores obtained by sampling with replacement on the prediction indices (n = 1000).

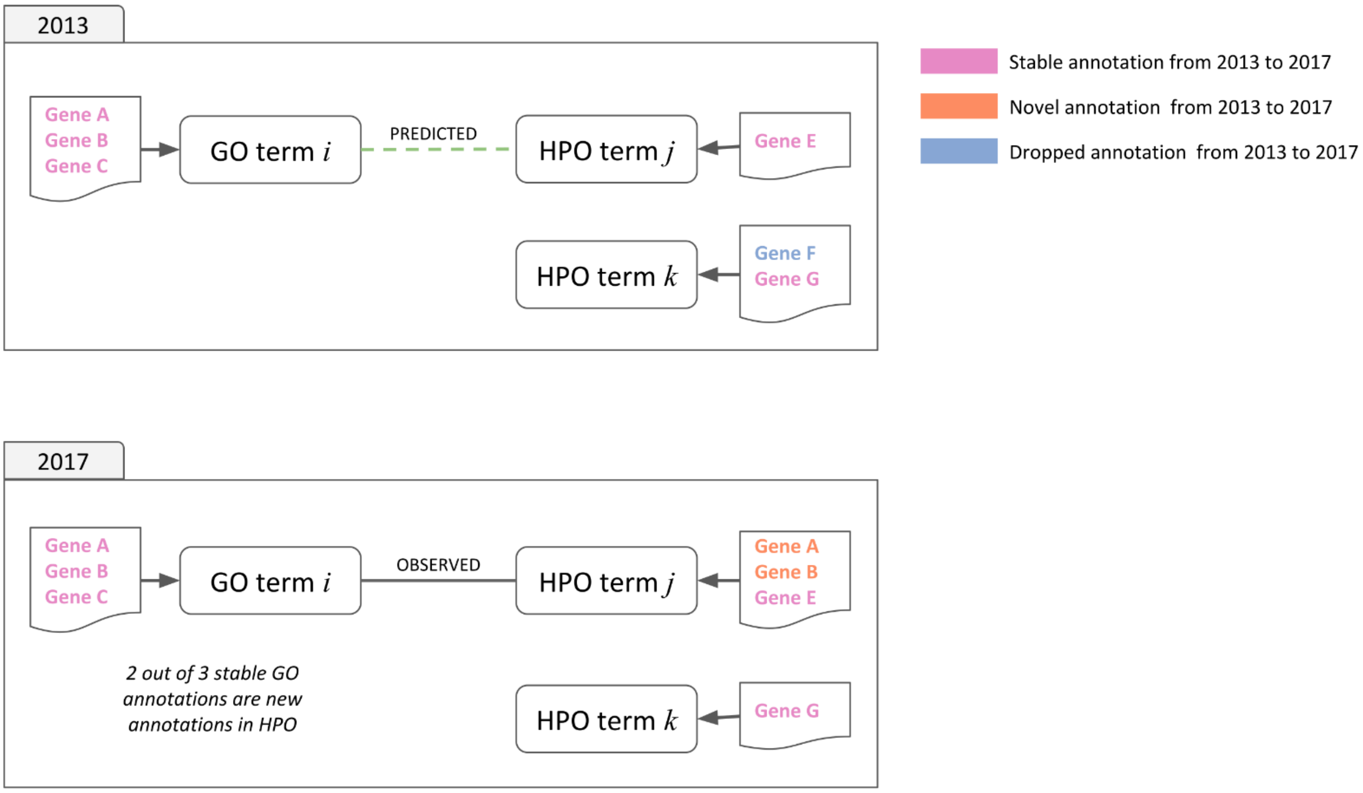


**Supplementary Figure 3.** Comparison of annotations in 2013 and 2017 ontology releases: stable annotations (pink), novel annotations (orange), dropped annotations (blue). In this example, genes A and B acquired an annotation to HPO term *j* in 2017 that, using 2013 releases, we predicted to have a high likelihood to be connected to GO term *i*.

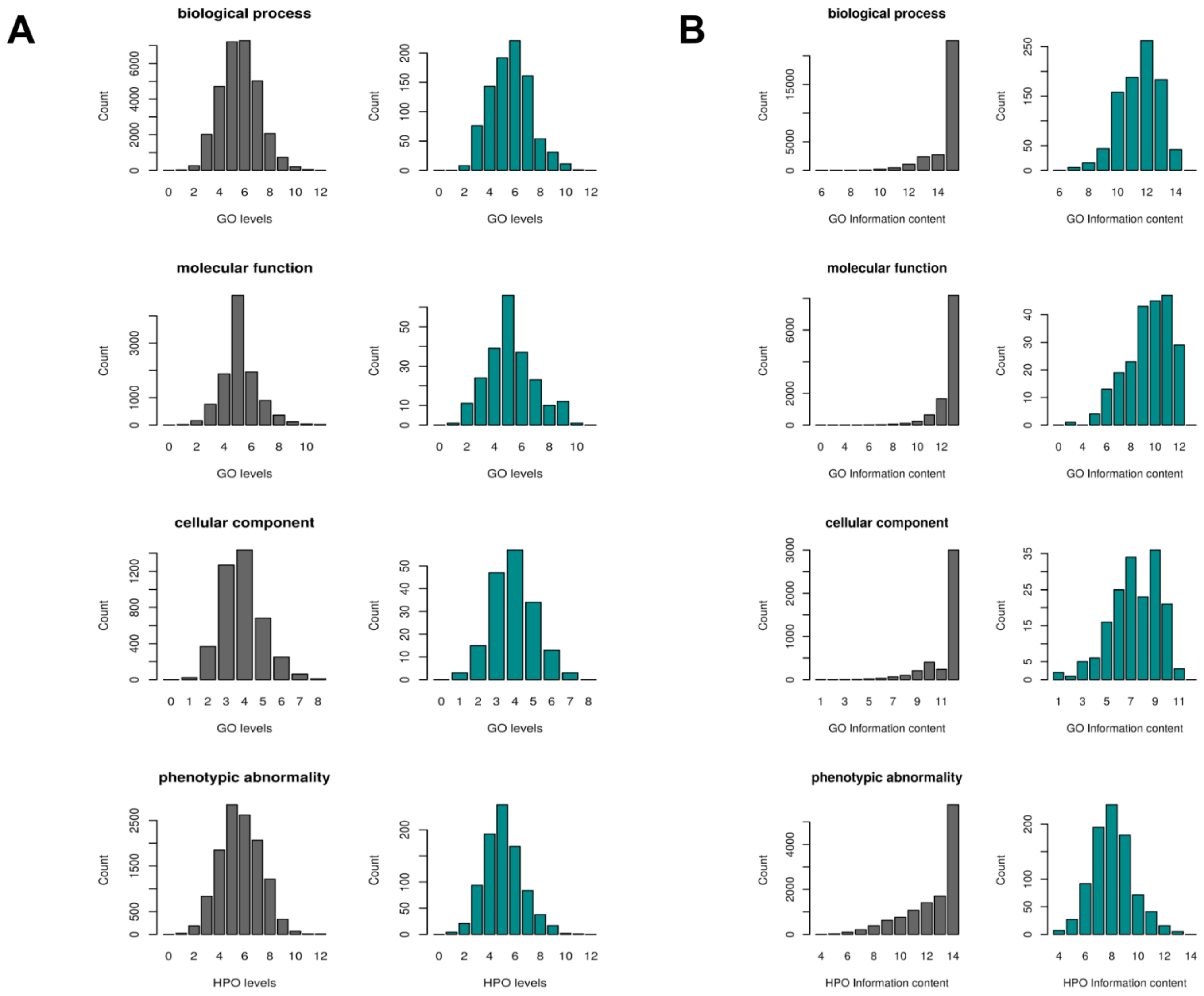


**Supplementary Figure 4.** [A] Distribution of graph levels of GO and HPO ontologies (gray) and terms in our predictions (green). [B] Distribution of Information content of GO and HPO ontologies (gray) and terms in our predictions (green).

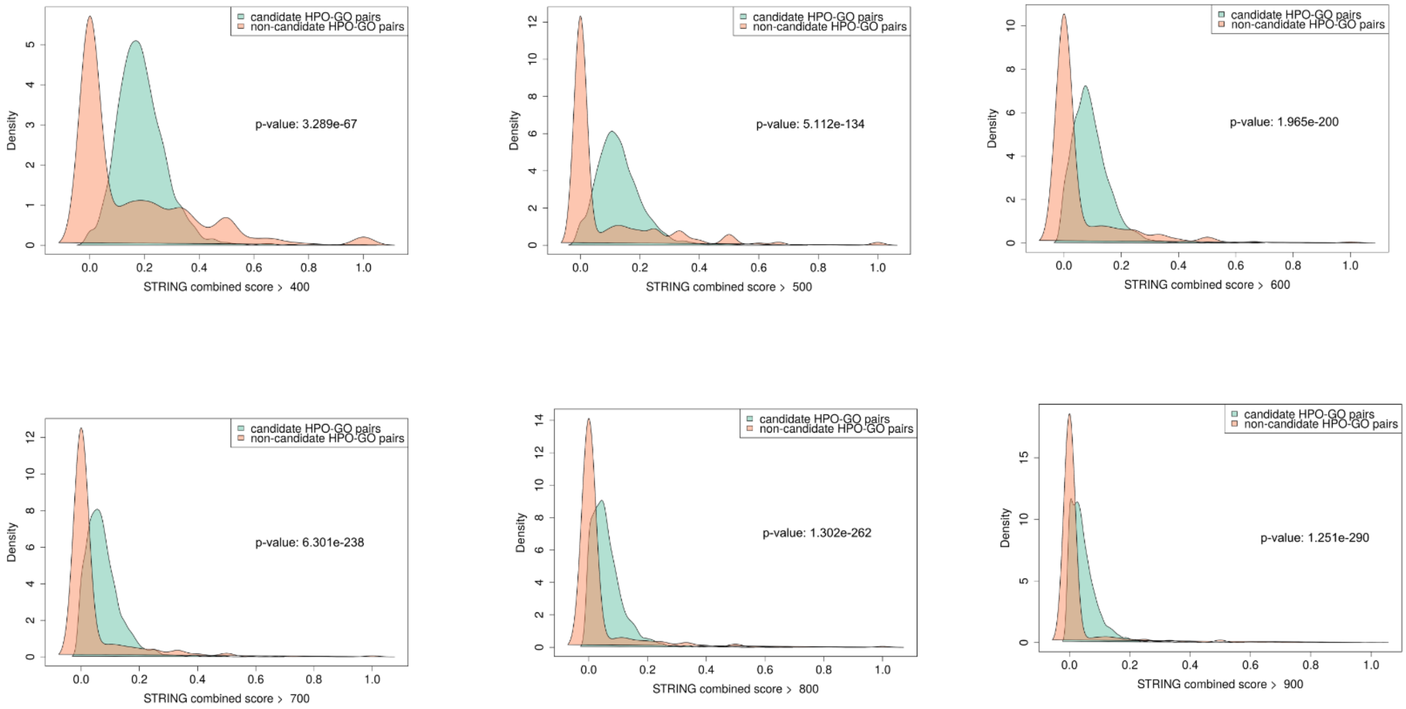


**Supplementary Figure 5.** Distributions of fraction of co-occurring genes annotated to predicted HPO-GO pairs found in STRING using increasing cutoff of STRING combined score (from 400, low confidence, to 900, high confidence). Significance of the difference between candidate and non-candidate HPO-GO terms was assessed by two-sided Wilcoxon signed-rank test.

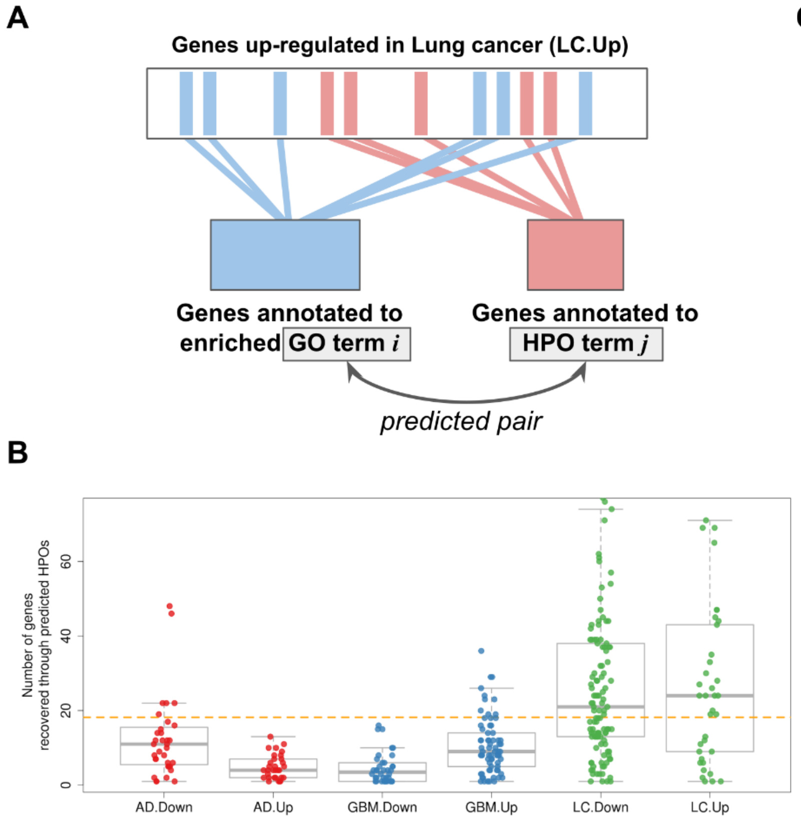


C

| | **GBM.Down** (N: 79) | **GBM.Up** (N: 204) |
|---|---|---|
| **AD.Down** (N: 202) | 21 (p-value 3.142e-10) | 1 (p-value 0.999) |
| **AD.Up** (N: 54) | 0 (p-value 0.720) | 13 (p-value 1.833e-06 ) |

| | **LC.Down** (N: 533) | **LC.Up** (N: 424) |
|---|---|---|
| **AD.Down** (N: 202) | 32 (p-value: 0.416) | 38 (p-value: 0.002) |
| **AD.Up** (N: 54) | 22 (p-value: 1.641e-06 ) | 6 (p-value: 0.513) |

| | **GBM.Down** (N: 79) | **GBM.Up** (N: 204) |
|---|---|---|
| **LC.Down** (N: 533) | 19 (p-value: 0.016) | 50 (p-value: 0.0002) |
| **LC.Up** (N: 424) | 10 (p-value: 0.391) | 44 (p-value: 4.929e-05) |

**Supplementary Figure 6.** Expanding candidate genes from results of functional enrichment analysis. [A] Genes annotated to an over-represented GO term (blue) are coupled with others drawn from the same gene set (up-regulated in Lung Cancer, LC.Up, in the example) and annotated to the corresponding predicted HPO. [B] Distributions of genes annotated to HPOs predicted to be associated with the enriched GO terms in the six gene sets. The average number is 18.19 (dashed horizontal orange line). [C] Tables reporting the number of genes recovered by predicted HPOs overlapping in the six gene sets (N, total number of genes annotated to predicted HPOs in the sets; p-values computed by hypergeometric test; the total number of genes annotated to HPOs is 3414, see Table 1). Green boxes indicate same directionality of expression profiles; orange boxes indicate inverse directionality of expression profiles. AD, Alzheimer's disease; LC, Lung Cancer; GBM, Glioblastoma; Up, up regulated; Down, down-regulated.

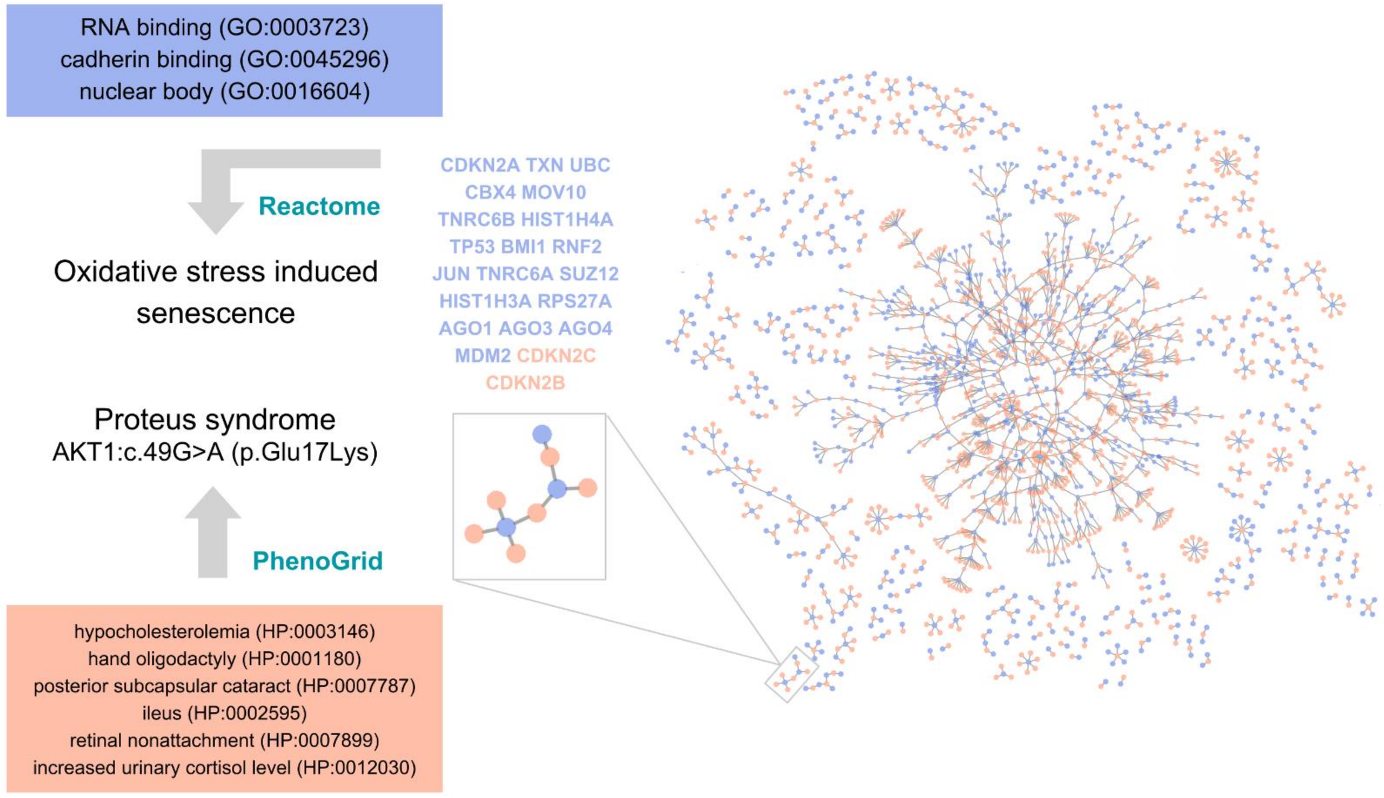


**Supplementary Figure 7.** Pathway analysis of clusters of candidate HPO-GO terms. A cluster (or a group of connected components) composed of 6 HPOs (orange dots) and 3 GOs (blue dots) is highlighted in the network of candidate HPO-GO terms (left). HPOs and GOs definitions, as well as genes corresponding annotated genes, are reported in orange and blue, respectively. Those genes are enriched in Reactome pathway oxidative stress induced senescence. Proteus syndrome, associated to a AKT1 variant inducing senescence events, is predicted by PhenoGrid based on those HPOs.

**Supplementary Tables**

| Ontology | Ontology version date | Sub-ontologies | Number of levels | Annotations | Annotations version date | Number of non-obsolete terms | Number of annotated genes |
|---|---|---|---|---|---|---|---|
| Gene Ontology (GO) | 29 June, 2013 | Biological process | 13 | GO Annotation (GOA) for Uniprot version 117 | 05 March, 2013 | 37,779 | 5,529 |
| | | Molecular function | 12 | | | | 7,139 |
| | | Cellular component | 9 | | | | 6,811 |
| | 23 May, 2017 | Biological process | 13 | GO Annotation (GOA) for Uniprot version 168 | 09 May, 2017 | 44,615 | 7,965 |
| | | Molecular function | 12 | | | | 11,309 |
| | | Cellular component | 9 | | | | 10,586 |
| Human Phenotype Ontology (HPO) | 31 May, 2013 | Phenotypic abnormality | 13 | HPO annotation files (see Methods) | 01 July, 2013 | 10,013 | 2,715 |
| | | Onset and clinical course | 5 | | | | |
| | | Mode of inheritance | 4 | | | | |
| | 13 April 2017 | Phenotypic abnormality | 13 | | 29 June, 2017 | 12,227 | 3,414 |
| | | Clinical modifier | 5 | | | | |
| | | Mode of inheritance | 4 | | | | |
| | | Mortality/Aging | 3 | | | | |
| | | Frequency | 2 | | | | |

**Supplementary Table 1**. Ontologies and annotations used in this work. Last column reports the number of human genes annotated with an experimental evidence code (see Methods).

| **Ontology release** | **HPO-GO term pairs** | **Path type** | **Average number of occurrences** | **Standard deviation of occurrences** |
|---|---|---|---|---|
| 2013 | disconnected | phghg | 6.33 | 32.57 |
| | | phphg | 23.32 | 125.59 |
| | | phggg | 0.09 | 0.86 |
| | | pphgg | 0.36 | 6.24 |
| | | ppphg | 0.09 | 0.82 |
| | | phgg | 0.03 | 0.46 |
| | | pphg | 0.05 | 0.47 |
| | connected | ppphg | 2.67 | 14.85 |
| | | pphgg | 1.77 | 12.76 |
| | | pphg | 0.76 | 6.17 |
| | | phghg | 375.73 | 1108.81 |
| | | phphg | 1671.42 | 5954.06 |
| | | phgg | 1.26 | 6.56 |
| | | phggg | 2.53 | 9.5 |
| 2017 | disconnected | phghg | 17.72 | 91.49 |
| | | phphg | 47.47 | 278.74 |
| | | phggg | 0.17 | 1.84 |
| | | phgg | 0.06 | 1 |
| | | ppphg | 0.08 | 0.77 |
| | | pphgg | 0.41 | 8.97 |
| | | pphg | 0.04 | 0.63 |
| | connected | phghg | 1063.35 | 3348.96 |
| | | phphg | 3511.57 | 12479.37 |
| | | ppphg | 3.17 | 18.5 |
| | | pphg | 0.97 | 7.75 |
| | | pphgg | 2.2 | 16.38 |
| | | phgg | 1.87 | 10.22 |
| | | phggg | 3.68 | 15.55 |

**Supplementary Table 2**. Type of paths and related statistics, for disconnected and connected HPO-GO term pairs randomly sampled from 2013 and 2017 ontology releases. Average number of paths per path type, and its standard deviation, are reported.

| Predictions using 2013 ontology releases | | | | | | | |
|---|---|---|---|---|---|---|---|
| Candidate HPO-GO term pairs | | | | Non-candidate HPO-GO term pairs | | | |
| HPO | GO | Likelihood of being connected | Likelihood of not being connected | HPO | GO | Likelihood of being connected | Likelihood of not being connected |
| HP:0002360 | GO:0006461 | 1.09E-05 | 1.02E-59 | HP:0010173 | GO:0044459 | 1.54E-05 | 9.24E-06 |
| HP:0100542 | GO:0000978 | 1.09E-05 | 1.11E-47 | HP:0006159 | GO:0035993 | 1.54E-05 | 9.24E-06 |
| HP:0002208 | GO:0018105 | 1.09E-05 | 7.20E-65 | HP:0010803 | GO:0051010 | 1.54E-05 | 9.24E-06 |
| HP:0002014 | GO:0048146 | 1.09E-05 | 2.32E-63 | HP:0200024 | GO:0009378 | 1.54E-05 | 9.24E-06 |
| HP:0001945 | GO:0051260 | 1.09E-05 | 2.88E-37 | HP:0009715 | GO:0005902 | 1.54E-05 | 9.24E-06 |
| HP:0000613 | GO:0051262 | 1.09E-05 | 6.42E-52 | HP:0000113 | GO:0006706 | 1.54E-05 | 9.24E-06 |
| HP:0001425 | GO:0009314 | 1.09E-05 | 1.76E-69 | HP:0010759 | GO:0086091 | 1.54E-05 | 9.24E-06 |
| HP:0100761 | GO:0030335 | 1.08E-05 | 4.74E-66 | HP:0011834 | GO:0043498 | 1.54E-05 | 9.24E-06 |
| HP:0002647 | GO:0001934 | 1.08E-05 | 9.75E-53 | HP:0004283 | GO:0043568 | 1.54E-05 | 9.24E-06 |
| HP:0000962 | GO:0004714 | 1.08E-05 | 5.40E-50 | HP:0001929 | GO:0004402 | 1.54E-05 | 9.24E-06 |
| HP:0001641 | GO:0000187 | 1.08E-05 | 6.69E-62 | HP:0011387 | GO:0030301 | 1.54E-05 | 9.24E-06 |
| HP:0003042 | GO:0043406 | 1.08E-05 | 1.38E-122 | HP:0000961 | GO:0010949 | 1.54E-05 | 9.24E-06 |
| HP:0001177 | GO:0008283 | 1.08E-05 | 1.14E-53 | HP:0006692 | GO:0005776 | 1.54E-05 | 9.24E-06 |
| HP:0001328 | GO:0000987 | 1.08E-05 | 7.07E-84 | HP:0010234 | GO:0043547 | 1.54E-05 | 9.24E-06 |
| HP:0004348 | GO:0005874 | 1.08E-05 | 2.74E-81 | HP:0003199 | GO:0042524 | 1.54E-05 | 9.24E-06 |
| HP:0001263 | GO:0033137 | 1.08E-05 | 5.67E-84 | HP:0001103 | GO:0005086 | 1.54E-05 | 9.24E-06 |
| HP:0003312 | GO:0001933 | 1.08E-05 | 6.03E-44 | HP:0002046 | GO:0060433 | 1.54E-05 | 9.24E-06 |
| HP:0000324 | GO:0030178 | 1.08E-05 | 6.50E-48 | HP:0006279 | GO:0006298 | 1.54E-05 | 9.24E-06 |
| HP:0001270 | GO:0006260 | 1.08E-05 | 4.91E-76 | HP:0100024 | GO:0043568 | 1.54E-05 | 9.24E-06 |
| HP:0002167 | GO:0030500 | 1.08E-05 | 8.37E-83 | HP:0007601 | GO:0035315 | 1.54E-05 | 9.24E-06 |
| HP:0001770 | GO:0007243 | 1.08E-05 | 5.39E-50 | HP:0002883 | GO:0032137 | 1.54E-05 | 9.24E-06 |
| HP:0002714 | GO:0043406 | 1.08E-05 | 1.89E-87 | HP:0003121 | GO:0043666 | 1.54E-05 | 9.24E-06 |
| HP:0000083 | GO:0030522 | 1.08E-05 | 4.56E-57 | HP:0011379 | GO:0043498 | 1.54E-05 | 9.24E-06 |
| HP:0000639 | GO:0031982 | 1.08E-05 | 6.89E-81 | HP:0001788 | GO:0035327 | 1.54E-05 | 9.24E-06 |
| HP:0009800 | GO:0005794 | 1.07E-05 | 3.32E-66 | HP:0002225 | GO:1900180 | 1.54E-05 | 9.24E-06 |
| HP:0000580 | GO:0005874 | 1.07E-05 | 3.91E-105 | HP:0006279 | GO:0043922 | 1.54E-05 | 9.24E-06 |
| HP:0001724 | GO:0005783 | 1.07E-05 | 2.06E-99 | HP:0002247 | GO:0046898 | 1.54E-05 | 9.24E-06 |
| HP:0000007 | GO:0046877 | 1.07E-05 | 9.56E-110 | HP:0000891 | GO:0005546 | 1.54E-05 | 9.24E-06 |
| HP:0000864 | GO:0001933 | 1.07E-05 | 1.14E-35 | HP:0007015 | GO:0043627 | 1.54E-05 | 9.24E-06 |
| HP:0000405 | GO:0008104 | 1.07E-05 | 2.51E-110 | HP:0100539 | GO:0032495 | 1.54E-05 | 9.24E-06 |
| HP:0001679 | GO:0034097 | 1.07E-05 | 2.51E-37 | HP:0001531 | GO:0000309 | 1.54E-05 | 9.24E-06 |
| HP:0000230 | GO:0007165 | 1.07E-05 | 1.06E-42 | HP:0008721 | GO:2001033 | 1.54E-05 | 9.24E-06 |
| HP:0001163 | GO:0097190 | 1.07E-05 | 6.68E-39 | HP:0001245 | GO:0003289 | 1.54E-05 | 9.24E-06 |

| | | | | | | | |
|---|---|---|---|---|---|---|---|
| HP:0002002 | GO:0045669 | 1.07E-05 | 1.31E-46 | HP:0100261 | GO:0046403 | 1.54E-05 | 9.24E-06 |
| HP:0001872 | GO:0043235 | 1.07E-05 | 1.35E-27 | HP:0009796 | GO:0060413 | 1.54E-05 | 9.24E-06 |
| HP:0100031 | GO:0001934 | 1.07E-05 | 1.14E-60 | HP:0005543 | GO:0004722 | 1.54E-05 | 9.24E-06 |
| HP:0000263 | GO:0005794 | 1.07E-05 | 6.57E-45 | HP:0000418 | GO:0006402 | 1.54E-05 | 9.24E-06 |
| HP:0000256 | GO:0043547 | 1.07E-05 | 1.44E-45 | HP:0003049 | GO:0032911 | 1.54E-05 | 9.24E-06 |
| HP:0008572 | GO:0008094 | 1.07E-05 | 1.78E-122 | HP:0008223 | GO:0004677 | 1.54E-05 | 9.24E-06 |
| HP:0000938 | GO:0016323 | 1.07E-05 | 2.85E-102 | HP:0000092 | GO:0060836 | 1.54E-05 | 9.24E-06 |
| HP:0001943 | GO:0010862 | 1.07E-05 | 2.57E-93 | HP:0012024 | GO:0008104 | 1.54E-05 | 9.24E-06 |
| HP:0001324 | GO:0045787 | 1.07E-05 | 8.77E-49 | HP:0007874 | GO:0071347 | 1.54E-05 | 9.24E-06 |
| HP:0000276 | GO:0000976 | 1.07E-05 | 1.01E-96 | HP:0006216 | GO:0033674 | 1.54E-05 | 9.24E-06 |
| HP:0002648 | GO:0032870 | 1.07E-05 | 2.61E-23 | HP:0011001 | GO:0015747 | 1.54E-05 | 9.24E-06 |
| HP:0000252 | GO:0035988 | 1.07E-05 | 2.94E-69 | HP:0002732 | GO:0045821 | 1.54E-05 | 9.24E-06 |
| HP:0007759 | GO:0003705 | 1.07E-05 | 3.90E-41 | HP:0004425 | GO:0008236 | 1.54E-05 | 9.24E-06 |
| HP:0100533 | GO:0048661 | 1.07E-05 | 7.04E-31 | HP:0007617 | GO:0007030 | 1.54E-05 | 9.24E-06 |
| HP:0001596 | GO:0042995 | 1.07E-05 | 2.32E-40 | HP:0100255 | GO:0008047 | 1.54E-05 | 9.24E-06 |
| HP:0003220 | GO:0030509 | 1.07E-05 | 1.15E-39 | HP:0000993 | GO:0007411 | 1.54E-05 | 9.24E-06 |
| HP:0000772 | GO:0030522 | 1.07E-05 | 1.49E-26 | HP:0004319 | GO:0060349 | 1.54E-05 | 9.24E-06 |
| HP:0001425 | GO:0051353 | 1.07E-05 | 1.13E-50 | HP:0000547 | GO:0042347 | 1.54E-05 | 9.24E-06 |
| HP:0000998 | GO:0006513 | 1.07E-05 | 2.36E-58 | HP:0011266 | GO:0001917 | 1.54E-05 | 9.24E-06 |
| HP:0000348 | GO:0015030 | 1.07E-05 | 1.63E-82 | HP:0001493 | GO:0030240 | 1.54E-05 | 9.24E-06 |
| HP:0001831 | GO:0000187 | 1.07E-05 | 1.31E-122 | HP:0001889 | GO:0010894 | 1.54E-05 | 9.24E-06 |
| HP:0000154 | GO:0071013 | 1.07E-05 | 1.34E-59 | HP:0002946 | GO:0010008 | 1.54E-05 | 9.24E-06 |
| HP:0001608 | GO:0009611 | 1.07E-05 | 2.75E-115 | HP:0007018 | GO:2001213 | 1.54E-05 | 9.24E-06 |
| HP:0008551 | GO:0001102 | 1.07E-05 | 3.13E-49 | HP:0000946 | GO:0046403 | 1.54E-05 | 9.24E-06 |
| HP:0100721 | GO:0043065 | 1.07E-05 | 1.47E-64 | HP:0002870 | GO:0030061 | 1.54E-05 | 9.24E-06 |
| HP:0009466 | GO:0034097 | 1.07E-05 | 1.69E-27 | HP:0007414 | GO:0050957 | 1.54E-05 | 9.24E-06 |
| HP:0006101 | GO:0000983 | 1.07E-05 | 5.51E-38 | HP:0009110 | GO:0034454 | 1.54E-05 | 9.24E-06 |
| HP:0002205 | GO:0043589 | 1.07E-05 | 3.39E-59 | HP:0002986 | GO:0071467 | 1.54E-05 | 9.24E-06 |
| HP:0002894 | GO:0006283 | 1.07E-05 | 1.45E-44 | HP:0001090 | GO:0004515 | 1.54E-05 | 9.24E-06 |
| HP:0000003 | GO:0070936 | 1.07E-05 | 1.98E-92 | HP:0001491 | GO:0020037 | 1.54E-05 | 9.24E-06 |
| HP:0002751 | GO:0001938 | 1.07E-05 | 6.52E-96 | HP:0004099 | GO:0030049 | 1.54E-05 | 9.24E-06 |
| HP:0100716 | GO:0006468 | 1.07E-05 | 7.59E-72 | HP:0005622 | GO:0032460 | 1.54E-05 | 9.24E-06 |
| HP:0006703 | GO:0043406 | 1.07E-05 | 2.99E-69 | HP:0001952 | GO:0060828 | 1.54E-05 | 9.24E-06 |
| HP:0000248 | GO:0035108 | 1.07E-05 | 5.78E-32 | HP:0009832 | GO:0045088 | 1.54E-05 | 9.24E-06 |
| HP:0004443 | GO:0005794 | 1.07E-05 | 8.63E-118 | HP:0006335 | GO:0060038 | 1.54E-05 | 9.24E-06 |
| HP:0002013 | GO:0030509 | 1.06E-05 | 5.27E-104 | HP:0000893 | GO:0045910 | 1.54E-05 | 9.24E-06 |
| HP:0001425 | GO:0071480 | 1.06E-05 | 2.57E-40 | HP:0002333 | GO:0042104 | 1.54E-05 | 9.24E-06 |
| HP:0004322 | GO:0043028 | 1.06E-05 | 2.81E-60 | HP:0001662 | GO:0008967 | 1.54E-05 | 9.24E-06 |

| | | | | | | | |
|---|---|---|---|---|---|---|---|
| HP:0000076 | GO:0018105 | 1.06E-05 | 3.74E-105 | HP:0002886 | GO:0034097 | 1.54E-05 | 9.24E-06 |
| HP:0003196 | GO:0045111 | 1.06E-05 | 3.57E-98 | HP:0003001 | GO:0004675 | 1.54E-05 | 9.24E-06 |
| HP:0000508 | GO:0071893 | 1.06E-05 | 1.10E-118 | HP:0000060 | GO:0045095 | 1.54E-05 | 9.24E-06 |
| HP:0002093 | GO:0016192 | 1.06E-05 | 2.10E-107 | HP:0006335 | GO:0051795 | 1.54E-05 | 9.24E-06 |
| HP:0007703 | GO:0015031 | 1.06E-05 | 2.88E-76 | HP:0002090 | GO:0022405 | 1.54E-05 | 9.24E-06 |
| HP:0002650 | GO:0009913 | 1.06E-05 | 9.75E-84 | HP:0012023 | GO:0043982 | 1.54E-05 | 9.24E-06 |
| HP:0002664 | GO:0070936 | 1.06E-05 | 2.46E-32 | HP:0001605 | GO:0061045 | 1.54E-05 | 9.24E-06 |
| HP:0000174 | GO:0060394 | 1.06E-05 | 3.33E-50 | HP:0001269 | GO:0000019 | 1.54E-05 | 9.24E-06 |
| HP:0100627 | GO:0070207 | 1.06E-05 | 3.88E-79 | HP:0100555 | GO:0052106 | 1.54E-05 | 9.24E-06 |
| HP:0001518 | GO:0001938 | 1.06E-05 | 5.24E-82 | HP:0010446 | GO:0004867 | 1.54E-05 | 9.24E-06 |
| HP:0001629 | GO:0043568 | 1.06E-05 | 4.38E-47 | HP:0000445 | GO:0033699 | 1.54E-05 | 9.24E-06 |
| HP:0000939 | GO:0006461 | 1.06E-05 | 3.26E-82 | HP:0003126 | GO:0005776 | 1.54E-05 | 9.24E-06 |
| HP:0000767 | GO:0003684 | 1.06E-05 | 5.40E-90 | HP:0000883 | GO:0048024 | 1.54E-05 | 9.24E-06 |
| HP:0001376 | GO:0043001 | 1.06E-05 | 4.96E-65 | HP:0002793 | GO:2000066 | 1.54E-05 | 9.24E-06 |
| HP:0001265 | GO:0010225 | 1.06E-05 | 1.72E-50 | HP:0001748 | GO:0006266 | 1.54E-05 | 9.24E-06 |
| HP:0000609 | GO:0005794 | 1.06E-05 | 4.01E-91 | HP:0100813 | GO:0022408 | 1.54E-05 | 9.24E-06 |
| HP:0001263 | GO:0006417 | 1.06E-05 | 1.42E-49 | HP:0003781 | GO:0010165 | 1.54E-05 | 9.24E-06 |
| HP:0002967 | GO:0010862 | 1.06E-05 | 4.66E-48 | HP:0002217 | GO:0030026 | 1.54E-05 | 9.24E-06 |
| HP:0002240 | GO:0010596 | 1.06E-05 | 3.52E-31 | HP:0100764 | GO:0007431 | 1.54E-05 | 9.24E-06 |
| HP:0000062 | GO:0007165 | 1.06E-05 | 1.40E-124 | HP:0010295 | GO:0015651 | 1.54E-05 | 9.24E-06 |
| HP:0004306 | GO:0043410 | 1.06E-05 | 7.93E-28 | HP:0000883 | GO:0004325 | 1.54E-05 | 9.24E-06 |
| HP:0000164 | GO:0004879 | 1.06E-05 | 7.33E-46 | HP:0200044 | GO:0030893 | 1.54E-05 | 9.24E-06 |
| HP:0000623 | GO:0005794 | 1.06E-05 | 1.19E-42 | HP:0010761 | GO:0034372 | 1.54E-05 | 9.24E-06 |
| HP:0007902 | GO:0000122 | 1.06E-05 | 1.41E-52 | HP:0000720 | GO:0055003 | 1.54E-05 | 9.24E-06 |
| HP:0000964 | GO:0009611 | 1.06E-05 | 1.55E-58 | HP:0100696 | GO:0015886 | 1.54E-05 | 9.24E-06 |
| HP:0001399 | GO:0097191 | 1.06E-05 | 3.00E-28 | HP:0009072 | GO:0030057 | 1.54E-05 | 9.24E-06 |
| HP:0000486 | GO:0030658 | 1.06E-05 | 4.34E-36 | HP:0005855 | GO:0072101 | 1.54E-05 | 9.24E-06 |
| HP:0000426 | GO:0000723 | 1.06E-05 | 6.39E-76 | HP:0004326 | GO:0043666 | 1.54E-05 | 9.24E-06 |
| HP:0001064 | GO:0044212 | 1.06E-05 | 6.99E-73 | HP:0002726 | GO:0070371 | 1.54E-05 | 9.24E-06 |

| Predictions using 2017 ontology releases | | | | | | | |
|---|---|---|---|---|---|---|---|
| Candidate HPO-GO term pairs | | | | Non-candidate HPO-GO term pairs | | | |
| HPO | GO | Likelihood of being connected | Likelihood of not being connected | HPO | GO | Likelihood of being connected | Likelihood of not being connected |
| HP:0001297 | GO:0033138 | 1.22E-05 | 1.07E-65 | HP:0001682 | GO:0010941 | 1.55E-05 | 1.05E-05 |
| HP:0000529 | GO:0000776 | 1.21E-05 | 2.32E-41 | HP:0007814 | GO:0006890 | 1.55E-05 | 1.05E-05 |
| HP:0000618 | GO:0045597 | 1.21E-05 | 1.65E-33 | HP:0010471 | GO:0042326 | 1.55E-05 | 1.05E-05 |

| | | | | | | | |
|---|---|---|---|---|---|---|---|
| HP:0000752 | GO:0000976 | 1.21E-05 | 2.11E-36 | HP:0001719 | GO:0003359 | 1.55E-05 | 1.05E-05 |
| HP:0004390 | GO:0010628 | 1.20E-05 | 3.55E-83 | HP:0002965 | GO:0005811 | 1.55E-05 | 1.05E-05 |
| HP:0000252 | GO:0001569 | 1.20E-05 | 1.44E-47 | HP:0011560 | GO:0005506 | 1.55E-05 | 1.05E-05 |
| HP:0005328 | GO:0043066 | 1.20E-05 | 1.34E-68 | HP:0003281 | GO:0042118 | 1.55E-05 | 1.05E-05 |
| HP:0003316 | GO:0010628 | 1.20E-05 | 3.25E-29 | HP:0001607 | GO:0010107 | 1.55E-05 | 1.05E-05 |
| HP:0012368 | GO:0048661 | 1.20E-05 | 9.65E-62 | HP:0003311 | GO:0000159 | 1.55E-05 | 1.05E-05 |
| HP:0000772 | GO:0033138 | 1.20E-05 | 8.04E-80 | HP:0012639 | GO:0005913 | 1.55E-05 | 1.05E-05 |
| HP:0200042 | GO:0003690 | 1.20E-05 | 8.92E-80 | HP:0002248 | GO:0042100 | 1.55E-05 | 1.05E-05 |
| HP:0001088 | GO:0044212 | 1.20E-05 | 2.68E-87 | HP:0100699 | GO:0035924 | 1.55E-05 | 1.05E-05 |
| HP:0009473 | GO:0005667 | 1.20E-05 | 2.73E-40 | HP:0005230 | GO:0045177 | 1.55E-05 | 1.05E-05 |
| HP:0000498 | GO:0001102 | 1.20E-05 | 6.79E-48 | HP:0000876 | GO:1903038 | 1.55E-05 | 1.05E-05 |
| HP:0100615 | GO:0004713 | 1.20E-05 | 3.27E-55 | HP:0002299 | GO:1900106 | 1.55E-05 | 1.05E-05 |
| HP:0000107 | GO:0034644 | 1.20E-05 | 2.71E-80 | HP:0012473 | GO:0032570 | 1.55E-05 | 1.05E-05 |
| HP:0002204 | GO:0000977 | 1.20E-05 | 3.67E-39 | HP:0100646 | GO:0046600 | 1.55E-05 | 1.05E-05 |
| HP:0001297 | GO:0016301 | 1.20E-05 | 5.33E-23 | HP:0010455 | GO:0000902 | 1.55E-05 | 1.05E-05 |
| HP:0001829 | GO:0001938 | 1.20E-05 | 1.33E-45 | HP:0007469 | GO:0016579 | 1.55E-05 | 1.05E-05 |
| HP:0000506 | GO:0043407 | 1.19E-05 | 7.06E-57 | HP:0006739 | GO:0042254 | 1.55E-05 | 1.05E-05 |
| HP:0000278 | GO:0016301 | 1.19E-05 | 8.82E-52 | HP:0001369 | GO:0006241 | 1.55E-05 | 1.05E-05 |
| HP:0003146 | GO:0003723 | 1.19E-05 | 3.30E-61 | HP:0002738 | GO:0001541 | 1.55E-05 | 1.05E-05 |
| HP:0001162 | GO:0045727 | 1.19E-05 | 1.27E-34 | HP:0011950 | GO:1904874 | 1.55E-05 | 1.05E-05 |
| HP:0000023 | GO:0043536 | 1.19E-05 | 1.48E-74 | HP:0200097 | GO:0015168 | 1.55E-05 | 1.05E-05 |
| HP:0002982 | GO:0031965 | 1.19E-05 | 3.20E-47 | HP:0001547 | GO:1904261 | 1.55E-05 | 1.05E-05 |
| HP:0001596 | GO:0045184 | 1.19E-05 | 3.04E-34 | HP:0000907 | GO:0001947 | 1.55E-05 | 1.05E-05 |
| HP:0001945 | GO:0072593 | 1.19E-05 | 2.71E-30 | HP:0009756 | GO:0016575 | 1.55E-05 | 1.05E-05 |
| HP:0001251 | GO:0038166 | 1.19E-05 | 2.33E-25 | HP:0005293 | GO:0032302 | 1.55E-05 | 1.05E-05 |
| HP:0003326 | GO:0006469 | 1.19E-05 | 2.32E-32 | HP:0012534 | GO:0070933 | 1.55E-05 | 1.05E-05 |
| HP:0004209 | GO:0048387 | 1.19E-05 | 5.46E-33 | HP:0008479 | GO:0017112 | 1.55E-05 | 1.05E-05 |
| HP:0000268 | GO:0045786 | 1.19E-05 | 1.07E-98 | HP:0002009 | GO:0071459 | 1.55E-05 | 1.05E-05 |
| HP:0004209 | GO:0016571 | 1.19E-05 | 7.63E-51 | HP:0004467 | GO:0004751 | 1.55E-05 | 1.05E-05 |
| HP:0001251 | GO:0044314 | 1.19E-05 | 2.18E-36 | HP:0001806 | GO:0071712 | 1.55E-05 | 1.05E-05 |
| HP:0000378 | GO:0001078 | 1.19E-05 | 5.43E-45 | HP:0002570 | GO:1901628 | 1.55E-05 | 1.05E-05 |
| HP:0000582 | GO:0043001 | 1.19E-05 | 4.72E-35 | HP:0001717 | GO:0007267 | 1.55E-05 | 1.05E-05 |
| HP:0005978 | GO:0000186 | 1.19E-05 | 1.47E-27 | HP:0004904 | GO:2000005 | 1.55E-05 | 1.05E-05 |
| HP:0000656 | GO:0000776 | 1.19E-05 | 1.65E-46 | HP:0100817 | GO:0005658 | 1.55E-05 | 1.05E-05 |
| HP:0011968 | GO:0006475 | 1.19E-05 | 1.45E-23 | HP:0100678 | GO:0036157 | 1.55E-05 | 1.05E-05 |
| HP:0012731 | GO:0010628 | 1.19E-05 | 5.58E-24 | HP:0002385 | GO:0004407 | 1.55E-05 | 1.05E-05 |
| HP:0003701 | GO:0055037 | 1.19E-05 | 1.01E-25 | HP:0000636 | GO:0045177 | 1.55E-05 | 1.05E-05 |
| HP:0000365 | GO:0051290 | 1.19E-05 | 6.34E-26 | HP:0002720 | GO:0006407 | 1.55E-05 | 1.05E-05 |

| | | | | | | | |
|---|---|---|---|---|---|---|---|
| HP:0000938 | GO:0007015 | 1.19E-05 | 2.26E-32 | HP:0001954 | GO:0006570 | 1.55E-05 | 1.05E-05 |
| HP:0002097 | GO:0005667 | 1.19E-05 | 8.46E-50 | HP:0002198 | GO:0000815 | 1.55E-05 | 1.05E-05 |
| HP:0000340 | GO:0043014 | 1.19E-05 | 1.76E-21 | HP:0002745 | GO:0042438 | 1.55E-05 | 1.05E-05 |
| HP:0000160 | GO:0001106 | 1.19E-05 | 2.15E-35 | HP:0007378 | GO:0007596 | 1.55E-05 | 1.05E-05 |
| HP:0002553 | GO:0000281 | 1.19E-05 | 3.08E-43 | HP:0000832 | GO:0043087 | 1.55E-05 | 1.05E-05 |
| HP:0000518 | GO:0033162 | 1.19E-05 | 3.19E-44 | HP:0008527 | GO:1905802 | 1.55E-05 | 1.05E-05 |
| HP:0000973 | GO:0008270 | 1.19E-05 | 2.37E-66 | HP:0002138 | GO:0042509 | 1.55E-05 | 1.05E-05 |
| HP:0007703 | GO:0060548 | 1.19E-05 | 6.69E-32 | HP:0002965 | GO:0030216 | 1.55E-05 | 1.05E-05 |
| HP:0000221 | GO:0043410 | 1.18E-05 | 6.10E-51 | HP:0000912 | GO:1905572 | 1.55E-05 | 1.05E-05 |
| HP:0000597 | GO:0045892 | 1.18E-05 | 4.43E-56 | HP:0004492 | GO:0061750 | 1.55E-05 | 1.05E-05 |
| HP:0000007 | GO:0010880 | 1.18E-05 | 2.11E-49 | HP:0007818 | GO:1902282 | 1.55E-05 | 1.05E-05 |
| HP:0000347 | GO:2000146 | 1.18E-05 | 9.46E-52 | HP:0010747 | GO:0016941 | 1.55E-05 | 1.05E-05 |
| HP:0008669 | GO:0042803 | 1.18E-05 | 5.09E-21 | HP:0005248 | GO:0010508 | 1.55E-05 | 1.05E-05 |
| HP:0000083 | GO:0036289 | 1.18E-05 | 2.38E-33 | HP:0010871 | GO:2000147 | 1.55E-05 | 1.05E-05 |
| HP:0008163 | GO:0010628 | 1.18E-05 | 3.01E-34 | HP:0000746 | GO:0043039 | 1.55E-05 | 1.05E-05 |
| HP:0000668 | GO:0031648 | 1.18E-05 | 1.80E-63 | HP:0000912 | GO:0032237 | 1.55E-05 | 1.05E-05 |
| HP:0002329 | GO:0000122 | 1.18E-05 | 5.38E-79 | HP:0000117 | GO:0007195 | 1.55E-05 | 1.05E-05 |
| HP:0002013 | GO:0046330 | 1.18E-05 | 3.51E-27 | HP:0004493 | GO:1904426 | 1.55E-05 | 1.05E-05 |
| HP:0000256 | GO:0005249 | 1.18E-05 | 2.58E-71 | HP:0000467 | GO:0040030 | 1.55E-05 | 1.05E-05 |
| HP:0000347 | GO:1904798 | 1.18E-05 | 9.27E-39 | HP:0002869 | GO:0032302 | 1.55E-05 | 1.05E-05 |
| HP:0000486 | GO:1990572 | 1.18E-05 | 1.08E-41 | HP:0004823 | GO:0046579 | 1.55E-05 | 1.05E-05 |
| HP:0000582 | GO:0048662 | 1.18E-05 | 3.56E-24 | HP:0001045 | GO:0004565 | 1.55E-05 | 1.05E-05 |
| HP:0011968 | GO:0097527 | 1.18E-05 | 1.77E-25 | HP:0006089 | GO:0005884 | 1.55E-05 | 1.05E-05 |
| HP:0000023 | GO:0090630 | 1.18E-05 | 3.53E-28 | HP:0002793 | GO:0005452 | 1.55E-05 | 1.05E-05 |
| HP:0000505 | GO:0008066 | 1.18E-05 | 7.96E-28 | HP:0004453 | GO:0005641 | 1.55E-05 | 1.05E-05 |
| HP:0001894 | GO:0051260 | 1.18E-05 | 1.77E-38 | HP:0003713 | GO:0009055 | 1.55E-05 | 1.05E-05 |
| HP:0001053 | GO:0097435 | 1.18E-05 | 2.03E-70 | HP:0004841 | GO:0030522 | 1.55E-05 | 1.05E-05 |
| HP:0008736 | GO:0010800 | 1.18E-05 | 6.63E-44 | HP:0010788 | GO:0005267 | 1.55E-05 | 1.05E-05 |
| HP:0000006 | GO:0033503 | 1.18E-05 | 1.78E-30 | HP:0003370 | GO:0030097 | 1.55E-05 | 1.05E-05 |
| HP:0001320 | GO:0007050 | 1.18E-05 | 5.66E-81 | HP:0003469 | GO:0032727 | 1.55E-05 | 1.05E-05 |
| HP:0000431 | GO:0000733 | 1.18E-05 | 1.24E-88 | HP:0001949 | GO:0000049 | 1.55E-05 | 1.05E-05 |
| HP:0000256 | GO:0055074 | 1.18E-05 | 5.58E-42 | HP:0001705 | GO:0032720 | 1.55E-05 | 1.05E-05 |
| HP:0000093 | GO:0006260 | 1.18E-05 | 5.78E-31 | HP:0000143 | GO:0005229 | 1.55E-05 | 1.05E-05 |
| HP:0001939 | GO:0001937 | 1.18E-05 | 2.88E-119 | HP:0003674 | GO:0046040 | 1.55E-05 | 1.05E-05 |
| HP:0008736 | GO:0008094 | 1.18E-05 | 2.66E-121 | HP:0100031 | GO:0005740 | 1.55E-05 | 1.05E-05 |
| HP:0001347 | GO:0043325 | 1.18E-05 | 7.75E-61 | HP:0000053 | GO:0055056 | 1.55E-05 | 1.05E-05 |
| HP:0001744 | GO:0071346 | 1.18E-05 | 1.27E-23 | HP:0008388 | GO:0002368 | 1.55E-05 | 1.05E-05 |
| HP:0001263 | GO:0030162 | 1.18E-05 | 1.51E-26 | HP:0007359 | GO:0048298 | 1.55E-05 | 1.05E-05 |

| | | | | | | | |
|---|---|---|---|---|---|---|---|
| HP:0001263 | GO:0034505 | 1.18E-05 | 7.56E-91 | HP:0000230 | GO:0050444 | 1.55E-05 | 1.05E-05 |
| HP:0002023 | GO:0001937 | 1.18E-05 | 1.41E-48 | HP:0006979 | GO:0070187 | 1.55E-05 | 1.05E-05 |
| HP:0000602 | GO:0034614 | 1.18E-05 | 1.33E-45 | HP:0000150 | GO:1904434 | 1.55E-05 | 1.05E-05 |
| HP:0000488 | GO:2000811 | 1.18E-05 | 8.76E-37 | HP:0001612 | GO:1900063 | 1.55E-05 | 1.05E-05 |
| HP:0002750 | GO:0098797 | 1.18E-05 | 5.71E-24 | HP:0002006 | GO:0001518 | 1.55E-05 | 1.05E-05 |
| HP:0000894 | GO:0003713 | 1.18E-05 | 2.47E-56 | HP:0003528 | GO:0046580 | 1.55E-05 | 1.05E-05 |
| HP:0000648 | GO:0001530 | 1.18E-05 | 1.66E-61 | HP:0100769 | GO:0031581 | 1.55E-05 | 1.05E-05 |
| HP:0000648 | GO:0001530 | 1.18E-05 | 1.66E-61 | HP:0003440 | GO:0061028 | 1.55E-05 | 1.05E-05 |
| HP:0000260 | GO:0048661 | 1.18E-05 | 5.84E-46 | HP:0000039 | GO:0010936 | 1.55E-05 | 1.05E-05 |
| HP:0006297 | GO:0045727 | 1.18E-05 | 2.72E-19 | HP:0006716 | GO:0036404 | 1.55E-05 | 1.05E-05 |
| HP:0001513 | GO:0090307 | 1.18E-05 | 8.03E-75 | HP:0002385 | GO:0033185 | 1.55E-05 | 1.05E-05 |
| HP:0000498 | GO:0042531 | 1.18E-05 | 6.81E-52 | HP:0004432 | GO:0036105 | 1.55E-05 | 1.05E-05 |
| HP:0000768 | GO:0005667 | 1.18E-05 | 3.78E-98 | HP:0003146 | GO:0001819 | 1.55E-05 | 1.05E-05 |
| HP:0100490 | GO:0008104 | 1.18E-05 | 7.56E-28 | HP:0004749 | GO:0043932 | 1.55E-05 | 1.05E-05 |
| HP:0000160 | GO:0035115 | 1.18E-05 | 2.10E-53 | HP:0003141 | GO:1990391 | 1.55E-05 | 1.05E-05 |
| HP:0002827 | GO:0000982 | 1.18E-05 | 2.79E-16 | HP:0007702 | GO:0008076 | 1.55E-05 | 1.05E-05 |
| HP:0000581 | GO:0034976 | 1.18E-05 | 1.41E-39 | HP:0002870 | GO:2000781 | 1.55E-05 | 1.05E-05 |
| HP:0002020 | GO:0001104 | 1.18E-05 | 8.87E-24 | HP:0000237 | GO:0003727 | 1.55E-05 | 1.05E-05 |
| HP:0001249 | GO:0030326 | 1.18E-05 | 5.55E-37 | HP:0009968 | GO:0032211 | 1.55E-05 | 1.05E-05 |
| HP:0000613 | GO:0042149 | 1.17E-05 | 1.05E-55 | HP:0001679 | GO:0030672 | 1.55E-05 | 1.05E-05 |
| HP:0000750 | GO:0006888 | 1.17E-05 | 2.11E-43 | HP:0000935 | GO:0005687 | 1.55E-05 | 1.05E-05 |

**Supplementary Table 3.** Top 100 (ascending rank order) candidate and non-candidate HPO-GO term pairs with corresponding likelihood of being connected and not connected inferred using ontology releases of 2013 and 2017.
